# Plantaricin JK from *Lactiplantibacillus plantarum* ZJ316 inhibits enterohemorrhagic Escherichia coli O104:H4 and alters fecal microbial communities *in vitro*

**DOI:** 10.64898/2026.10.20.760378

**Authors:** Shumin Wang, Jiaqian Yan, Bin Yu, Ning Ding, Xiaolu Li, Shuxin Wang, Jiahao Jiang, Xiaodan Zhao, Qing Gu, Ping Li

## Abstract

Enterohemorrhagic *Escherichia coli* (EHEC) O104:H4 is a highly virulent foodborne pathogen for which antibiotic therapy is contraindicated, severely restricting conventional treatment options; therefore, the development of novel antimicrobial strategies is essential. This study investigated the antibacterial activity and gut microbiota-modulating effects of plantaricin JK (PlnJK), a two-peptide bacteriocin from *Lactiplantibacillus plantarum*. Chemically synthesized PlnJK exhibited a minimum inhibitory concentration (MIC) of 32 μM against *E. coli* O104:H4, disrupting membrane integrity as evidenced by increased extracellular conductivity, ATP depletion, and ultrastructural damage. In an *in vitro* fecal fermentation model, PlnJK reduced *Escherichia-Shigella* abundance from 36.70% to 30.60% which contains *E. coli* O104:H4-related taxa, enriched *Clostridium* (5.70% vs 2.40% in infection group), *Rikenellaceae*, and *Lactobacillaceae*, and significantly elevated short-chain fatty acid (SCFA) levels including butyrate (p < 0.01) and isovalerate (p < 0.05). To translate this into a deliverable format, CRISPR-Cas9-engineered *L. plantarum* strains with plnJK knockout, complementation, and overexpression were constructed. Compared with the chemically synthesized peptide, LAB-derived PlnJK—especially from the overexpression strain—showed superior efficacy in restoring microbial diversity and enhancing SCFA production, significantly increased Shannon index and reduced dominance (p < 0.01), while promoting butyrate (p < 0.01) and isovalerate (p < 0.01) production compared to the knockout control. The engineered strains further revealed a clear dose-dependent effect, as overexpression potentiated while knockout abolished the beneficial modulation. These findings demonstrate that PlnJK acts through membrane disruption and beneficially modulates gut microecology, with LAB-mediated in situ delivery offering a promising strategy for anti-EHEC live biotherapeutic development.

**Importance:** *Enterohemorrhagic Escherichia coli* O104:H4 causes life-threatening hemorrhagic colitis and hemolytic uremic syndrome, yet antibiotic therapy is contraindicated because it induces Shiga toxin release, leaving few treatment options. This study shows that plantaricin JK (PlnJK), a class IIb two-peptide bacteriocin characterized mainly against Gram-positive bacteria, inhibits this Gram-negative pathogen at 32 µM by disrupting membrane integrity. In an *in vitro* fecal fermentation model, PlnJK suppressed *Escherichia – Shigella*, increased community diversity, and elevated butyrate and isovalerate production, indicating concurrent pathogen control and microecological recovery. Using CRISPR-Cas9-engineered *Lactiplantibacillus plantarum* strains carrying *plnJK* knockout, complementation, and overexpression, we establish a dose-dependent causal link between PlnJK production and these effects, and show that LAB-mediated *in situ* delivery outperforms the chemically synthesized peptide. These findings extend the known activity spectrum of PlnJK and support a food-grade, antibiotic-free strategy against EHEC.

## Introduction

*Escherichia coli* is a Gram-negative, rod-shaped, facultatively anaerobic bacterium. It is a common gut commensal; however, certain strains can acquire virulence genes and cause disease(Croxen et al., 2013; Kaper et al., 2004). Pathogenic *E. coli* strains are classified into different pathotypes based on clinical syndromes, among which enterohemorrhagic *E. coli* (EHEC) is one of the most concerning, with an estimated 265,000 infections annually in the United States(Croxen et al., 2013; Perraud & Sperandio, 2025). EHEC infection can progress from watery diarrhea to bloody diarrhea and may develop into hemolytic uremic syndrome (HUS), characterized by microangiopathic hemolytic anemia, thrombocytopenia, and acute renal failure(Eltorki et al., 2026; Navaneethan & Giannella, 2008). The 2011 German O104:H4 outbreak, which caused over 4,000 infections, more than 900 HUS cases, and 54 deaths, highlighted its high virulence, strong colonization, Shiga toxin 2 (Stx2) production, and multidrug resistance(Croxen et al., 2013). Conventional treatments such as plasmapheresis and corticosteroids lack clear benefit, and antibiotics are contraindicated because they induce Stx release(Ma et al., 2026; Menne et al., 2012). Moreover, recent studies demonstrate that antibiotics can induce massive Stx release from EHEC, and are thus contraindicated for EHEC infections(Ma et al., 2026). These limitations underscore the urgent need for novel antimicrobial strategies with high specificity, low resistance risk, and compatibility with the host microbiota, such as bacteriocins(Bhattarai et al., 2026; Mihailovskaya et al., 2023).

Bacteriocins are ribosomally synthesized antimicrobial peptides, divided into class I (post-translationally modified) and class II (unmodified or disulfide-bonded only). Unlike broad-spectrum antibiotics, they can precisely kill pathogens without disrupting the commensal microbiota. Nisin and pediocin PA-1 have been approved by the FDA for food preservation, validating their safety. Bacteriocins exhibit diverse mechanisms of action, including inhibition of cell wall synthesis, disruption of cell membranes, and interference with transcription and translation. Some bacteriocins, such as darobactin, are active against Gram-negative drug-resistant bacteria. They can also be engineered through mutagenesis and incorporation of non-natural amino acids to optimize their activity and stability(Heilbronner et al., 2021; Sugrue et al., 2024).

Plantaricin PlnJK is a class IIb two-peptide bacteriocin from *Lactiplantibacillus plantarum* C11. Anderssen et al. (1998) first demonstrated that an equimolar mixture of PlnJ and PlnK is required for nanomolar-level activity, whereas each peptide alone exhibits at least 1,000-fold lower activity, and that this synergy is strictly specific. PlnJK mainly inhibits Gram-positive bacteria closely related to the producer but is inactive against *Carnobacterium*(Anderssen et al., 1998). In the following year, Moll et al. (1999) confirmed that PlnJK forms anion-selective pores, dissipates the transmembrane pH gradient, and causes intracellular acidification and cell death. On this basis, Diep et al. (2009) provided a systematic genetic characterization, showing that *plnJ* and *plnK* are located adjacently in the *plnJKLR* operon, ensuring equimolar expression; their precursors carry a double-glycine leader that is cleaved by the ABC transporter PlnG/H. Mature PlnJ and PlnK are cationic peptides composed of 25 and 32 amino acids, respectively, and adopt amphiphilic α-helices in membrane-mimetic environments, a conformation essential for pore formation(Anderssen et al., 1998; Diep et al., 2009; Moll et al., 1999). However, these findings cannot be directly extrapolated to Gram-negative pathogens such as *E. coli* O104:H4, whose outer membrane presents an additional permeability barrier to many bacteriocins. Direct evaluation is therefore needed to determine the inhibitory activity of PlnJK against this pathogen. Furthermore, antibacterial activity observed in pure culture does not necessarily predict pathogen suppression within complex gut microbial communities or the effects of treatment on the local microbial and metabolic environment. Accordingly, this study evaluated the antibacterial activity of PlnJK against *E. coli* O104:H4 and assessed its effects on gut microbial composition and SCFA profiles.

Compared with traditional isolation and purification from culture supernatants of natural strains, chemical synthesis, particularly solid-phase peptide synthesis, can yield a single homogeneous component with purity as high as >90% or even 98%, with a defined composition and free of any bacterial metabolites. For this reason, the use of metagenomic analysis to predict novel bacteriocins, followed by chemical synthesis to validate their activity, has become a cutting-edge approach for discovering and identifying novel antimicrobial peptides(Zhang et al., 2025). However, chemical synthesis of bacteriocins also has notable limitations: high cost for long peptides, lack of post-translational modifications, and susceptibility to proteolytic degradation, resulting in poor oral bioavailability(Sadeeq et al., 2025; Soltani et al., 2021). Except for nisin, most bacteriocins have not yet obtained food regulatory approval, and chemical synthesis cannot mimic the sustained in situ release by natural producer strains in the gut(Sadeeq et al., 2025; Soltani et al., 2021). Therefore, practical application still requires integration with strategies such as heterologous expression, formulation encapsulation, or engineered bacterial delivery.

Lactic acid bacteria (LAB) have well-recognized food safety status and intestinal adaptability, and their bacteriocins can selectively eliminate pathogens, promote beneficial bacteria, and increase SCFA production, thereby exerting anti-inflammatory and gut barrier-protective effects(W. Guo et al., 2025; Ismael et al., 2024; Mejía-Caballero & Marco, 2026). Bacteriocin-producing LAB can serve as live biotherapeutics that persistently colonize the gut and exclude pathogens. CRISPR-Cas genome editing enables scarless gene knockout and site-specific modification in LAB without antibiotic markers, meeting food-grade engineering requirements; CRISPRi, in turn, allows reversible regulation of gene expression, providing powerful tools for elucidating the biological characteristics of PlnJK and improving strains(Ullah et al., 2025; Zhou et al., 2023). Thus, applying CRISPR-Cas technology to LAB chassis represents a highly effective strategy for systematically elucidating the biological characteristics of PlnJK and promoting its translational application.

Here, we first evaluated the *in vitro* antibacterial activity and mechanisms of chemically synthesized PlnJK against *E. coli* O104:H4 by measuring membrane permeability, intracellular ATP, and ultrastructural changes, and assessed its impact on gut microbiota and SCFA metabolism using an *in vitro* fecal fermentation model. We then constructed engineered *L. plantarum* strains with *plnJK* knockout, complemented expression, and overexpression to systematically evaluate the selective antibacterial and microecological regulatory effects of LAB-derived PlnJK during *E. coli* O104:H4 infection. This study aims to provide a theoretical basis and technical support for developing anti-EHEC live biotherapeutic products based on LAB-derived bacteriocins.

## 2. Materials and Methods

### 2.1 Preparation and Formulation of Bacteriocin PlnJK

PlnJ and PlnK peptides were synthesized by GL Biochem Ltd. (Shanghai, China) via solid-phase peptide synthesis (SPPS) and purified by reverse-phase high-performance liquid chromatography (RP-HPLC). The purified PlnJ (purity 98.29%) and PlnK (purity 98.81%) were stored as lyophilized powder at −80°C. Prior to use, the peptides were dissolved in 0.05% acetic acid buffer and mixed at an equimolar ratio to obtain the PlnJK working solution, which was freshly prepared before each experiment.

### 2.2 Determination of Minimum Inhibitory Concentration (MIC)

The MIC of PlnJK against *Escherichia coli* O104:H4 was determined using the broth microdilution method described by Wiegand et al.(Wiegand et al., 2008), with minor modifications. Different concentrations of PlnJK (1–512 μM) were co-incubated with exponential-phase *E. coli* O104:H4 (OD₆₀₀ = 0.2) for 12 h. Bacterial growth was assessed by measuring OD₆₀₀, and the MIC was defined as the lowest concentration of PlnJK that completely inhibited detectable bacterial growth under the assay conditions.

### 2.3 Measurement of Membrane Conductivity and Intracellular ATP Levels

Exponential-phase bacterial cells were harvested and resuspended in HEPES buffer. PlnJK was added at 1 × MIC, and changes in extracellular conductivity and intracellular ATP levels were monitored using a conductivity meter and an ATP assay kit (Beyotime), respectively. Treatment with 1% Triton X-100 served as the positive control, and 0.05% acetic acid served as the negative control.

### 2.4 Electron Microscopy Observation

Bacterial cells were treated with PlnJK at 1 × MIC for 3 h, followed by fixation, dehydration, and drying. Changes in cell surface morphology and internal ultrastructure were examined by scanning electron microscopy (SEM) and transmission electron microscopy (TEM), respectively.

### 2.5 *In Vitro* Gut Microbiota Simulation

#### 2.5.1 Fecal Sample Collection and Experimental Groups

Morning fecal samples were collected from five healthy volunteers (Hangzhou, aged 20–30 years, with no antibiotic use within the preceding three months), all of whom provided informed consent. Fecal slurries were prepared. The experimental groups were as follows: Ctrl (blank control), Ec (inoculated with *E. coli* O104:H4), and PlnJK (inoculated with *E. coli* O104:H4 + 32 μM PlnJK). Fermentation was carried out at 37°C for 24 h under anaerobic conditions.

#### 2.5.2 16S rRNA Sequencing

Total DNA was extracted from the fermented samples, and the V3-V4 hypervariable region was amplified using primers 338F (5′-ACTCCTACGGGAGGCAGCA-3′) and 806R (5′-GGACTACHVGGGTWTCTAAT-3′). Sequencing was performed on the Illumina platform. Alpha diversity indices (Chao1, Shannon, and Simpson), beta diversity (PCoA and NMDS), and differentially abundant taxa between groups (LEfSe, LDA > 3) were analyzed using QIIME.

#### 2.5.3 Short-Chain Fatty Acid (SCFA) Analysis

The concentrations of seven SCFAs, including acetate, propionate, and butyrate, were determined by gas chromatography–mass spectrometry (GC-MS).

### 2.6 Construction of plnJK Knockout and Derivative Strains

#### 2.6.1 Strains and Plasmids

*Lactiplantibacillus plantarum* ZJ316 was originally isolated from the feces of a healthy newborn and has been deposited in the China Center for Type Culture Collection (CCTCC) under accession number CCTCC NO: M 208077. Its complete genome sequence is available in the GenBank database under accession number CP004082. The knockout vector pLCNICK was purchased from Addgene, and the expression vector pNZ44 was kindly provided by Prof. Paul Ross, University College Cork, Ireland. *Escherichia coli* DH5α was used for plasmid propagation.

#### 2.6.2 Construction of the Knockout Vector

The CRISPR-Cas9-based method described by Song et al.(Song et al., 2017). was followed. Primers (Table 1) were designed to amplify the upstream and downstream homology arms of the *plnJK* gene as well as the gRNA scaffold + sgRNA. These fragments were assembled into a knockout cassette by overlap-extension PCR. The pLCNICK vector was double-digested with ApaI and XbaI, ligated with the knockout cassette, and transformed into *E. coli* DH5α. Transformants were selected on erythromycin-containing plates and verified by sequencing.

**Table 1.**
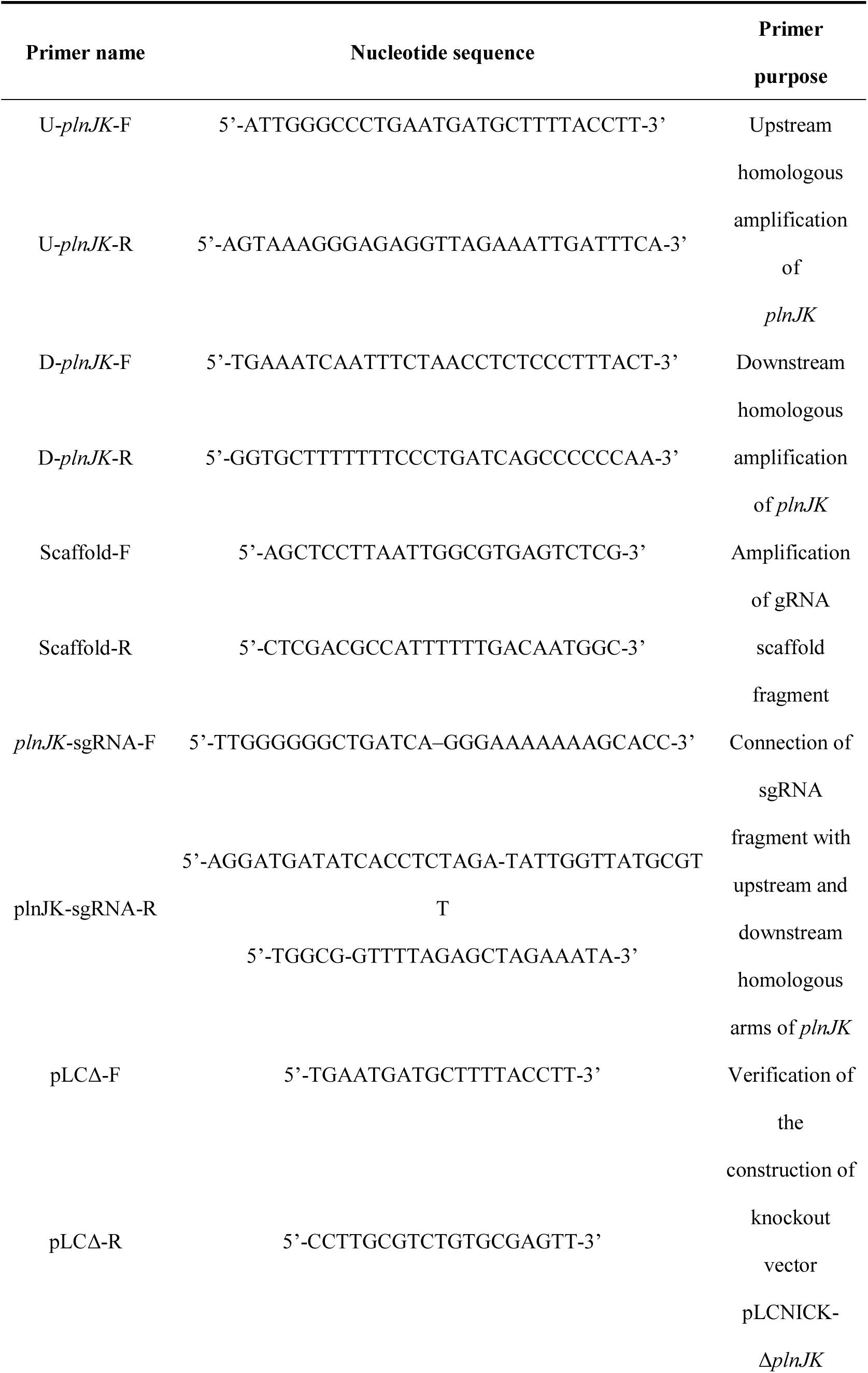
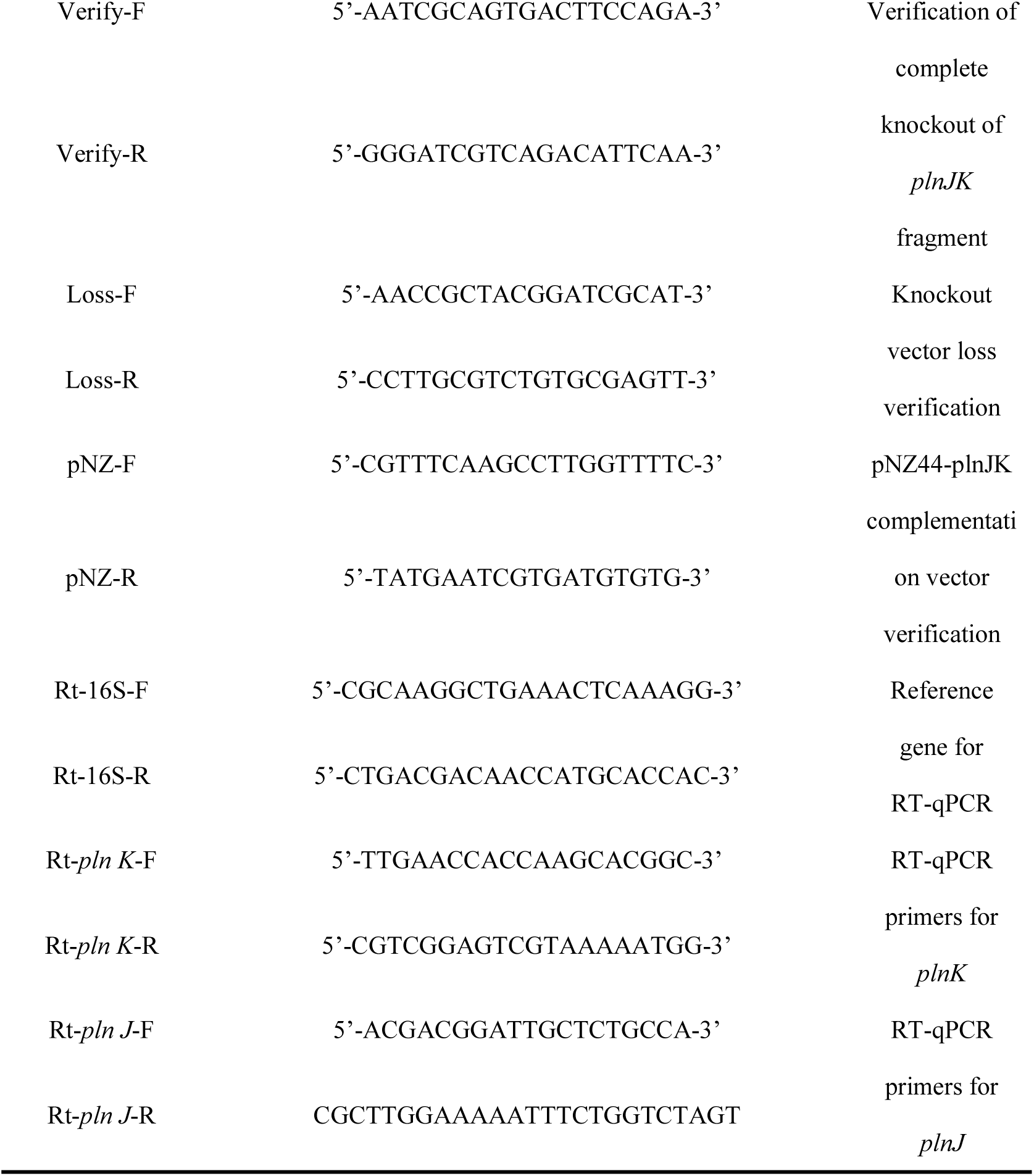
Primers used in the experiments.

#### 2.6.3 Electroporation and Screening of Knockout Strains

Competent cells of *L. plantarum* ZJ316 were prepared and electroporated with the knockout vector (2.5 kV). Positive clones were selected on MRS agar plates containing erythromycin, and *plnJK* deletion was confirmed by PCR and sequencing. The resulting strain was subjected to serial passage in antibiotic-free medium to cure the plasmid, yielding the markerless knockout strain ZJ316*ΔplnJK*.

#### 2.6.4 Construction of Complemented and Overexpression Strains

The full-length *plnJK* gene was amplified and cloned into pNZ44 (PstI/XbaI) to generate the expression vector pNZ44-*plnJK*. This construct was electroporated into ZJ316 and ZJ316Δ*plnJK* competent cells, and positive clones were selected on MRS agar plates containing chloramphenicol (30 μg/mL), yielding the overexpression strain ZJ316::*plnJK* and the complemented strain ZJ316Δ*plnJK*::*plnJK*, respectively.

#### 2.6.5 RT-qPCR Validation

Total RNA was extracted from the wild-type, knockout, complemented, and overexpression strains using a bacterial total RNA extraction kit (Tiangen, China). Following reverse transcription, the relative expression levels of *plnJ* and *plnK* were determined by RT-qPCR, with 16S rRNA serving as the internal reference.

#### 2.6.6 Growth Curve Analysis

Each strain was inoculated into MRS broth and cultured at 37°C for 24 h. OD₆₀₀ values were measured every 2 h to generate growth curves.

### 2.7 Modulation of Gut Microecology by LAB-Derived Plantaricin JK

#### 2.7.1 Tolerance to Simulated Gastrointestinal Fluids

Simulated gastric fluid (SGF) containing pepsin (pH 2.0, 3.0, and 4.0) and simulated intestinal fluid (SIF) containing pancreatin and bile salts at various concentrations (0.1%, 0.2%, and 0.4%) were prepared according to the method of Leeansyah et al.(Leeansyah et al., 2013). Cell suspensions of the four strains (ZJ316, ZJ316Δ*plnJK*, ZJ316Δ*plnJK*::*plnJK*, and ZJ316::*plnJK*) were inoculated into the above solutions and incubated at 37°C. Samples were collected at 0, 2, and 4 h, and viable cell counts were determined by plating on MRS agar. Tolerance was expressed as survival rate (viable cell count after treatment / initial viable cell count ×100%).

#### 2.7.2 *In Vitro* Gut Microbiota Simulation Experiment

Four additional groups were included based on the experimental setup described in Section 2.5.1: (4) WT group (infection + wild-type ZJ316), (5) KO group (infection + knockout strain ZJ316Δ*plnJK*), (6) RE group (infection + complemented strain ZJ316Δ*plnJK*::*plnJK*), and (7) Over group (infection + overexpression strain ZJ316::*plnJK*). Fecal sample collection, fermentation, 16S rRNA sequencing, and SCFA analysis followed Section 2.5.

### 2.8 Statistical Analysis

All experimental data are presented as mean ± standard deviation (Mean ± SD). Data were processed and analyzed using GraphPad Prism 9.4.1. One-way ANOVA followed by Tukey’s post hoc test was used for multiple-group comparisons. A p value < 0.05 was considered statistically significant.

## 3. Results

### 3.1 Effect of PlnJK against *Escherichia coli* O104:H4

#### 3.1.1 Inhibition of *Escherichia coli* O104:H4 growth by PlnJK

The antibacterial activity of Pln JK against Escherichia coli O104:H4 was determined using the broth microdilution method. As shown in Fig.1A, Pln JK inhibited the growth of E. coli O104:H4 in a concentration-dependent manner. The OD600 values gradually declined at concentrations of 1-16 μM, dropped sharply to near zero at 32 μM, and remained stable at higher concentrations. Therefore, the minimum inhibitory concentration (MIC) of Pln JK against *E. coli* O104:H4 was determined to be 32 μM.

**Figure 1.**
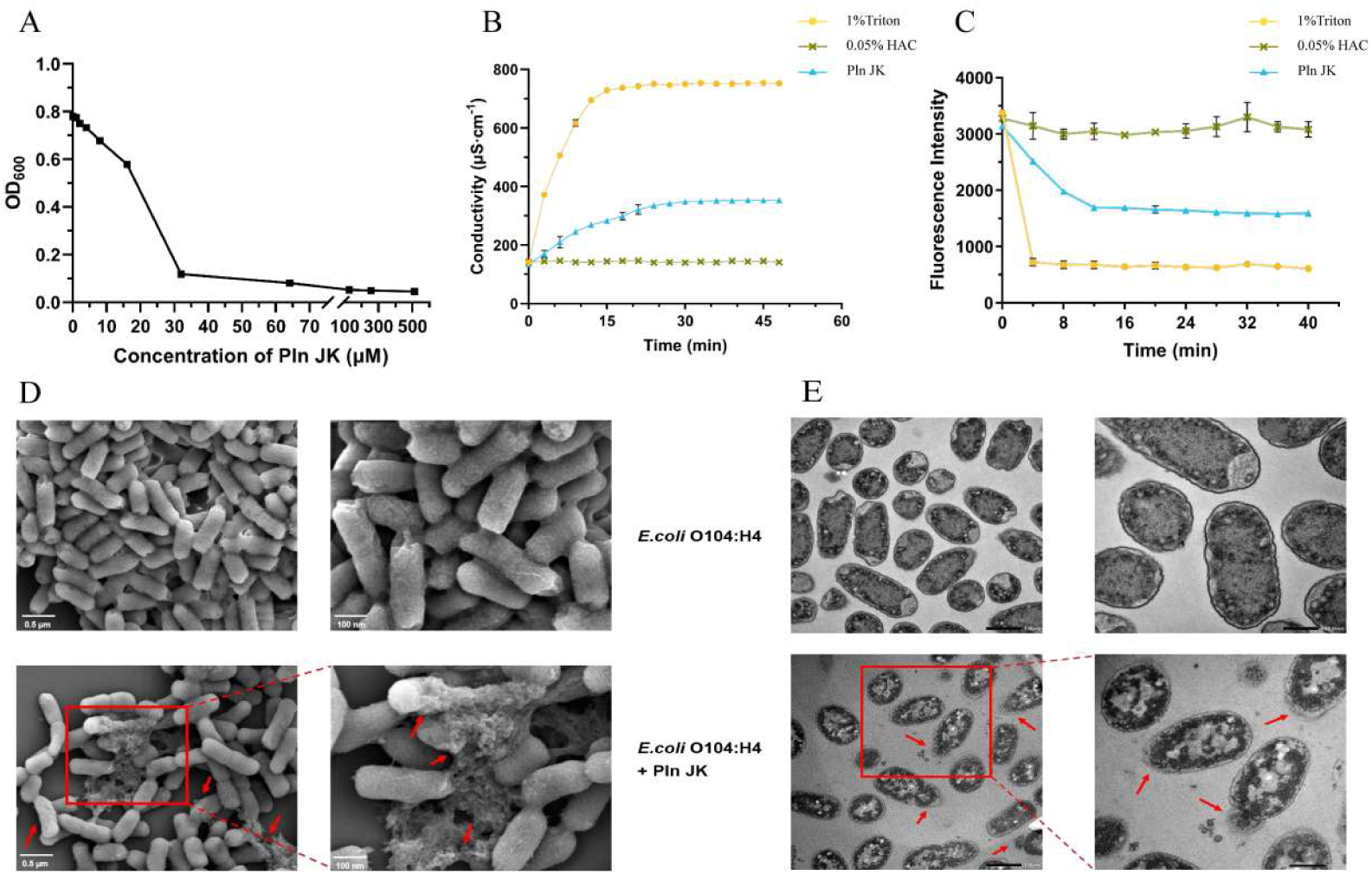
Antibacterial activity and membrane damage mechanisms of PlnJK against Escherichia coli O104:H4. (A) MIC determination (MIC = 32 μM). (B) Changes in extracellular conductivity. (C) Changes in intracellular ATP levels. (D) SEM images (scale bars: 0.5 μm, 100 nm). (E) TEM images (scale bars: 0.5 μm, 100 nm). Red arrows indicate membrane rupture (D) and plasmolysis (E).

#### 3.1.2 Effect of PlnJK on membrane permeability and intracellular ATP levels in *Escherichia coli* O104:H4

Membrane integrity disruption was assessed by measuring changes in extracellular conductivity. As shown in Fig.1B, the conductivity of the positive control group (1% Triton X-100) increased rapidly within 1-12 min and then stabilized, whereas the negative control group (0.05% acetic acid buffer) exhibited negligible change throughout the monitoring period. Upon addition of PlnJK at a final concentration of 32 μM, the conductivity of the bacterial suspension showed a marked and sustained increase, reaching approximately 348.00 μS/cm at around 30 min and leveling off thereafter, indicating leakage of intracellular ions. As shown in Fig.1C, compared with the negative control, intracellular ATP levels in *E. coli* O104:H4 decreased sharply within 4 min of PlnJK treatment and dropped to a low, stable level after 16 min. These results are consistent with the observed increase in conductivity and indicate that PlnJK increased membrane permeability, leading to rapid depletion of intracellular ATP.

#### 3.1.3 Effect of PlnJK on the ultrastructure of Escherichia coli O104:H4 cells

The surface morphology of *E. coli* O104:H4 cells before and after PlnJK treatment was examined by SEM. As shown in Fig. 1D, untreated control cells appeared plump with smooth surfaces and a typical short-rod shape. In contrast, after treatment with 32 μM PlnJK for 3 h, cells exhibited morphological alterations, including rough, collapsed surfaces and, in some cases, severe membrane rupture with leakage of intracellular contents. TEM further revealed ultrastructural changes. As shown in Fig.1E, untreated *E. coli* O104:H4 cells displayed intact and tightly adherent cell membrane and wall, with evenly distributed cytoplasmic contents. Following PlnJK treatment, marked plasmolysis was observed, along with uneven cytoplasmic distribution, vacuolation, damaged membrane integrity, and extensive leakage of cellular contents.

### 3.2 Effect of PlnJK on *in vitro* simulated gut microbiota

#### 3.2.1 Effect of PlnJK on gut microbiota alpha diversity

The alpha diversity of gut microbiota in different treatment groups was analyzed by 16S rRNA high-throughput sequencing. As shown in Fig.2A, compared with the control group (Ctrl), the *E. coli* O104:H4 infection group (Ec) exhibited significantly reduced Chao1 and Shannon indices (p < 0.01, p < 0.05), indicating decreased richness and diversity. In contrast, the PlnJK-treated group (PlnJK) showed recovered Chao1 and Shannon indices relative to the Ec group. Consistent trends in Dominance and Simpson indices further indicated that PlnJK could partially mitigate the gut microbiota dysbiosis caused by *E. coli* O104:H4.

**Figure 2.**
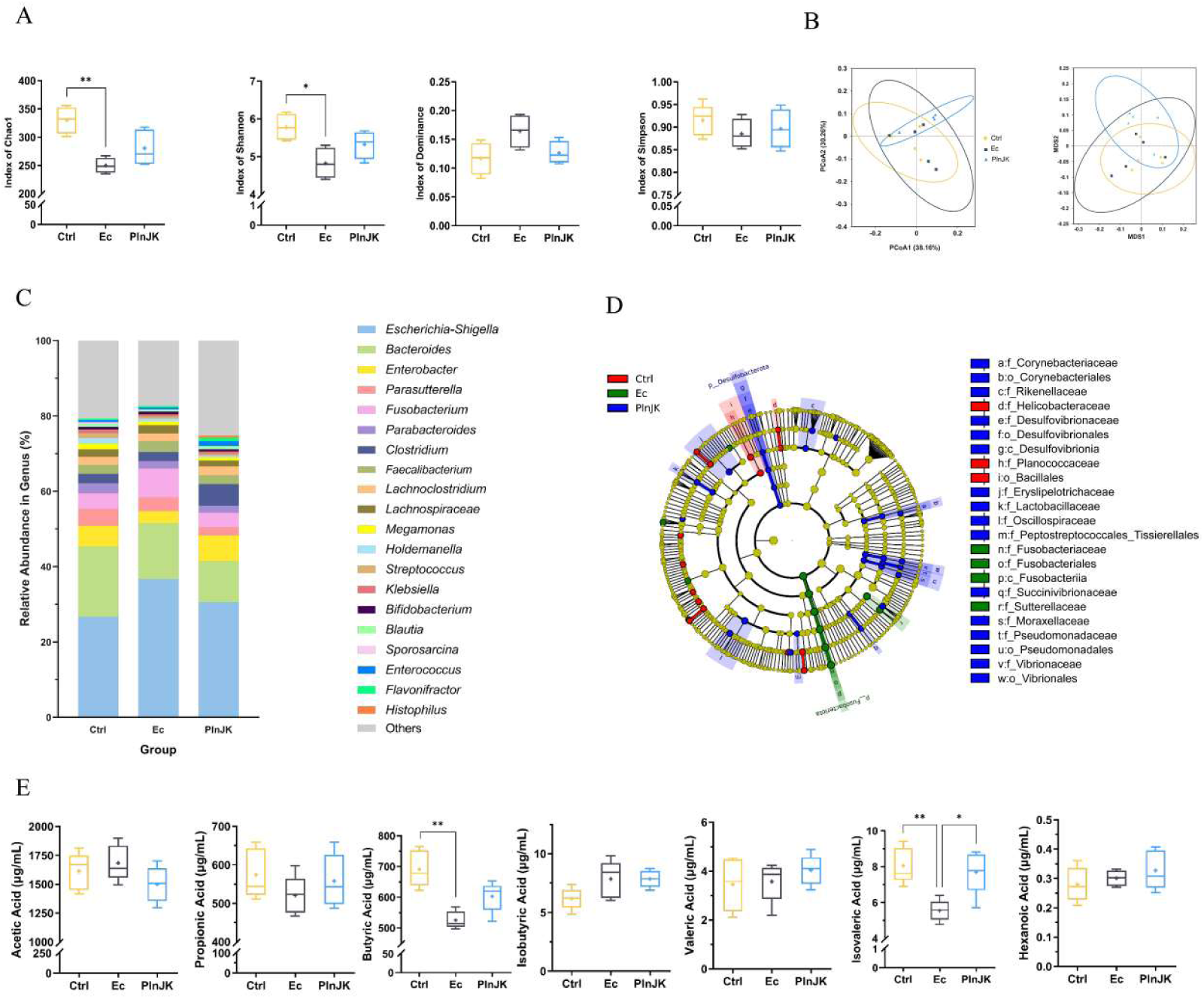
Effects of PlnJK on *in vitro* gut microbiota structure and metabolism. **(A) Alpha diversity indices.** (B) Beta diversity (PCoA, NMDS). (C) Relative abundance at genus level (top 20). (D) LEfSe differential taxa (LDA > 3). (E) SCFA concentrations. Ctrl, blank control; Ec, O104:H4 infection; PlnJK, infection + PlnJK. *p < 0.05, **p < 0.01.

#### 3.2.2 Effect of PlnJK on gut microbiota beta diversity

PCoA and NMDS based on Weighted UniFrac distance were used to assess differences in community structure among groups. As shown in Fig.2B, in both ordination plots, samples from the Ctrl and Ec groups clustered relatively close with within-group dispersion, whereas the PlnJK group samples were clearly separated from the Ec group and more tightly clustered. This indicates that PlnJK altered the microbial community structure caused by *E. coli* O104:H4, leading to a significant shift in the overall gut microbiota composition.

#### 3.2.3 Effect of PlnJK on gut microbiota composition

The relative abundance of gut microbiota was analyzed at the genus level. As shown in Fig.2C, in the Ec group, the relative abundance of *Escherichia-Shigella* increased markedly from 26.70% (Ctrl) to 36.70%, becoming the dominant taxon. After PlnJK treatment, its abundance decreased to 30.60%, indicating effective inhibition of *E. coli* O104:H4 by PlnJK. In addition, the relative abundance of Clostridium was significantly higher in the PlnJK group (5.70%) than in the Ctrl (2.50%) and Ec (2.40%) groups; many members of this genus are known to be beneficial SCFA producers.

#### 3.2.4 Differential analysis of gut microbiota structure by LEfSe

LEfSe analysis (LDA > 3) was used to identify differentially abundant microbial taxa among groups. As shown in Fig.2D, the PlnJK group was enriched in the most differential taxa, mainly including *Corynebacteriaceae, Rikenellaceae, Oscillospiraceae, and Lactobacillaceae.* Previous studies have shown that increased abundance of *Rikenellaceae* is associated with a reduced risk of inflammatory bowel disease, and *Oscillospiraceae* and *Lactobacillaceae* are known to play positive roles in maintaining intestinal microecological balance. These results suggest that PlnJK not only inhibits pathogenic bacteria but also promotes the growth of beneficial microbes in the gut.

#### 3.2.5 Effect of PlnJK on gut short-chain fatty acid levels

The concentrations of SCFAs in fermented samples were determined by GC-MS. As shown in Fig.2E, compared with the Ec group, the PlnJK group showed increased levels of propionate, butyrate, valerate, isovalerate, and hexanoate to varying degrees. Notably, isovalerate was significantly elevated (p < 0.05), and butyrate showed a highly significant increase (p < 0.01). SCFAs, particularly butyrate and isovalerate, are key microbial metabolites that play critical roles in maintaining intestinal barrier integrity, exerting anti-inflammatory effects, and regulating immunity. These findings further validate that PlnJK has the probiotic function of enriching gut microbiota and modulating microbial metabolites.

### 3.3 Construction of the *plnJK* knockout vector in *Lactiplantibacillus plantarum* ZJ316 and mutant validation

#### 3.3.1 Construction of the knockout vector pLCNICK-Δ*plnJK*

To construct a vector for targeted knockout of the *plnJK* gene, the upstream and downstream homology arms (U-*plnJK* and D-*plnJK*) were first amplified by PCR. Agarose gel electrophoresis showed that both fragments were approximately 1000 bp, matching the expected sizes. Overlap-extension PCR (SOE-PCR) was then used to fuse the two homology arms, yielding the U-*plnJK*-D fragment (∼2000 bp). Meanwhile, the gRNA scaffold fragment (∼379 bp) was amplified from the pLCNICK plasmid, and the sgRNA targeting *plnJK* was inserted via nested PCR, generating the gRNA scaffold + sgRNA fragment (∼141 bp). The U-*plnJK*-D fragment and the gRNA scaffold + sgRNA fragment were further fused by SOE-PCR to produce the final knockout template U-*plnJK*-D + gRNA scaffold + sgRNA (∼2090 bp) (Fig.3A). The pLCNICK plasmid was linearized by double digestion with ApaI and XbaI, and the large backbone (∼12,000 bp) was recovered. The knockout template was ligated into the linearized vector and transformed into *E. coli* DH5α competent cells. Colony PCR of 24 single colonies identified 5 clones with a specific band of approximately 2368 bp, confirming successful construction of the knockout vector pLCNICK-Δ*plnJK* (Fig.3B).

**Figure 3.**
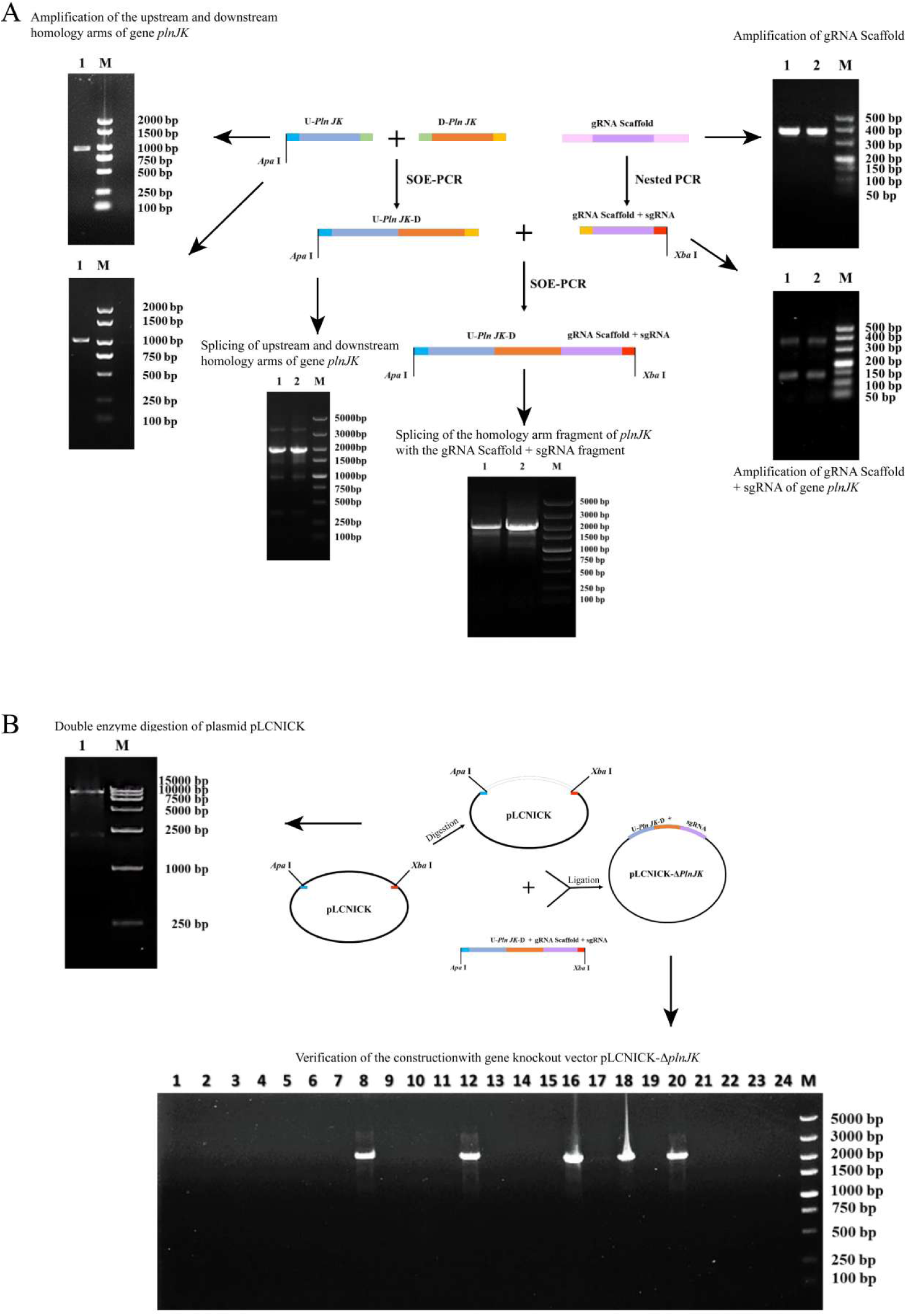
Construction strategy and PCR verification of the *plnJK* knockout vector pLCNICK-Δ*plnJK*. A) The upstream and downstream homology arms of the target gene *plnJK* (U-*plnJK* and D-*plnJK*, both ∼1000 bp) and the gRNA scaffold (∼379 bp) were amplified by PCR, with a specific sgRNA introduced. These fragments were sequentially assembled via SOE-PCR to generate the linear knockout template U-*plnJK*-D-gRNA scaffold-sgRNA (∼2090 bp). Lane numbers : different amplified homology arms or assembled fragments; M, DNA molecular weight marker.(B) The pLCNICK plasmid was double-digested with ApaI and XbaI to obtain the linearized backbone (∼12,000 bp), which was ligated with the knockout template from (A) and transformed into *E. coli* DH5α. Twenty-four single colonies were picked for colony PCR screening.

#### 3.3.2 Construction of the *plnJK* knockout mutant

The constructed knockout vector pLCNICK-Δ*plnJK* was introduced into *L. plantarum* ZJ316 competent cells by electroporation. Colony PCR screening using primers flanking the *plnJK* gene showed that positive clones yielded an amplicon of approximately 400 bp, whereas wild-type *L. plantarum* ZJ316 produced a band of ∼750 bp, confirming successful deletion of the target gene (Fig.4A). Sequencing of the PCR product further verified that it contained only the upstream and downstream homology arms without the *plnJK* gene sequence, thus confirming the knockout (Fig. 3.4B). The positive clone was subsequently subjected to serial passage in antibiotic-free MRS medium to cure the vector. PCR identification using the original vector as a control showed no amplification of vector-specific bands in the passaged strain, demonstrating the generation of the plasmid-free knockout mutant *L. plantarum* ZJ316Δ*plnJK* (Fig.4A).

**Figure 4.**
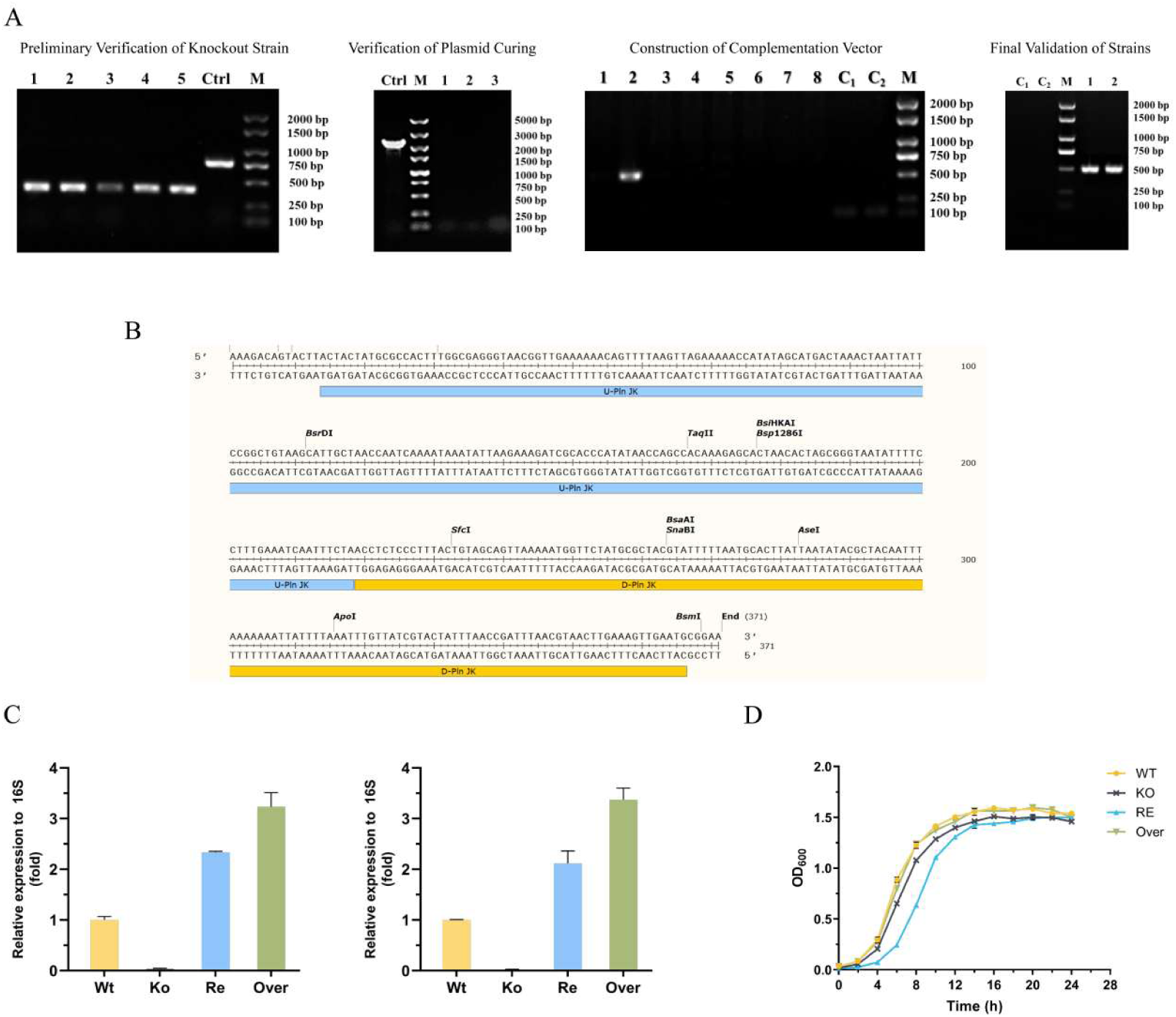
Construction and verification of plnJK knockout, complemented, and overexpression strains. (A) Colony PCR verification and plasmid curing identification. Left: preliminary verification of knockout strains (1–5, transformants; Ctrl, wild-type L. plantarum ZJ316). Middle: plasmid curing verification (Ctrl, uncured strain; 1–3, cured strains). Right: verification of complemented expression vector construction, with transformants showing an amplicon of ∼500 bp.(B) Sequencing verification of the knockout region, showing only the upstream and downstream homology arm sequences.(C) RT-qPCR analysis of relative expression levels of plnJ (left) and plnK (right) in different strains.(D) Growth curves of the four strains.

#### 3.3.3 Construction of complemented and overexpression strains

To further verify the function of the *plnJK* gene, we constructed the complemented expression vector pNZ44-*plnJK*. The *plnJK* gene fragment (372 bp) was ligated into pNZ44 and transformed into *E. coli* DH5α. Colony PCR showed that positive clones produced an amplicon of approximately 500 bp, whereas the empty vector control yielded only a ∼100 bp band, confirming successful plasmid construction (Fig.4A). The recombinant plasmid was electroporated into *L. plantarum* ZJ316Δ*plnJK* and wild-type *L. plantarum* ZJ316, respectively, generating the complemented mutant ZJ316Δ*plnJK*::*plnJK* and the overexpression mutant ZJ316::*plnJK*. PCR identification confirmed that both strains produced the expected ∼500 bp band, while the control strains without the plasmid showed no amplification, indicating successful introduction of the vector (Fig.4A).

#### 3.3.4 Analysis of target gene expression and strain growth characteristics

The relative expression levels of *plnJ* and *plnK* in different strains were quantified by RT-qPCR. The results showed that *plnJK* transcripts were nearly undetectable in the knockout strain *L. plantarum* ZJ316Δ*plnJK*, whereas the complemented strain ZJ316Δ*plnJK*::*plnJK* and the overexpression strain ZJ316::*plnJK* exhibited significantly higher expression of both genes compared to the wild-type *L. plantarum* ZJ316 (Fig.4C). Furthermore, growth curve analysis revealed that all four strains displayed similar growth trends, entering logarithmic phase at 4 h and reaching stationary phase at approximately 14 h, indicating that neither *plnJK* deletion nor introduction of the expression vector significantly affected normal growth (Fig.4D).

### 3.4 Modulatory effects of LAB-derived bacteriocin PlnJK on gut microecology

#### 3.4.1 Effect of strain-derived PlnJK on gut microbiota diversity

We first evaluated the alpha diversity of gut microbiota across different treatment groups. As shown in Fig.5A, the Ace index showed an increasing trend in the wild-type *L. plantarum* ZJ316-treated group (WT), the complemented group (RE), and the overexpression group (Over) compared with the Ec and KO groups. Notably, for the Shannon index (Fig.5A, top right), the Over group exhibited a significant increase compared with both the Ec and KO groups. Conversely, the Simpson and Dominance indices (Fig.5A, bottom) exhibited decreasing trends in the WT, RE, and Over groups. Specifically, the Over group showed a significantly lower Dominance index compared with the Ec and KO groups (p < 0.05 and p < 0.01). These results indicate that highly expressed PlnJK significantly increased the richness and evenness of the gut microbiota. Beta diversity analysis further revealed overall structural differences. As shown in Figure.5B, PCoA and NMDS based on Weighted UniFrac distance showed that samples from the WT, RE, and Over groups clustered tightly and separated distinctly from the Ec and KO groups. This indicates that *L. plantarum* ZJ316 and its produced PlnJK significantly altered the community structure disrupted by *E. coli* O104:H4, promoting the recovery of gut microecology.

**Figure 5.**
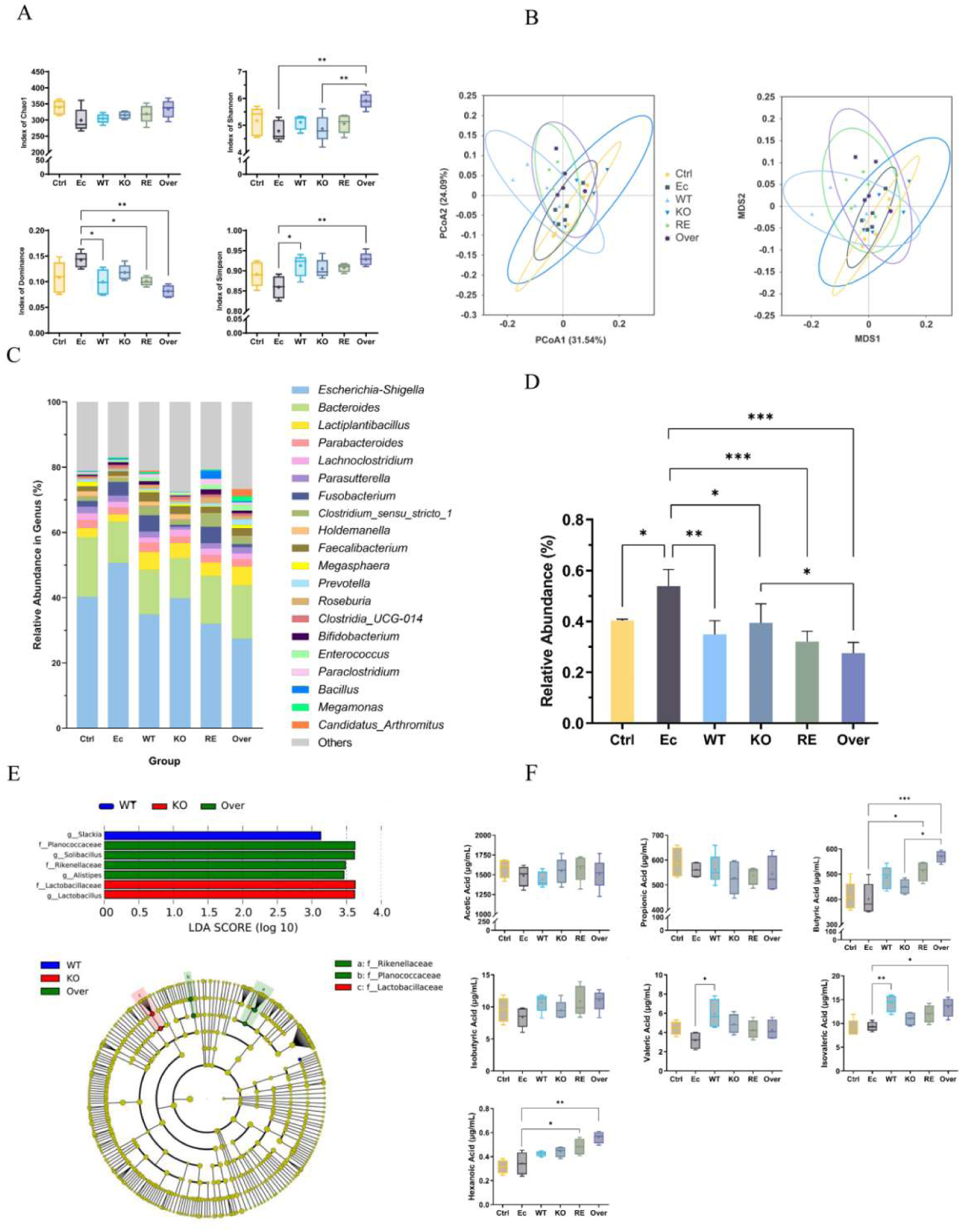
Modulatory effects of strain-derived PlnJK on gut microecology. (A) Alpha diversity indices (Chao1, Shannon, Simpson, and Dominance) across groups.(B) Beta diversity analysis based on Weighted UniFrac distances (PCoA and NMDS).(C) Relative abundance of gut microbiota at the genus level (top 20).(D) Relative abundance of the genus Escherichia-Shigella.(E) LEfSe analysis of differentially abundant taxa among WT, KO, and Over groups (LDA > 3).(F) Concentrations of short-chain fatty acids (SCFAs), including butyrate, isobutyrate, isovalerate, and valerate.Ctrl, blank control; Ec, O104:H4 infection; WT, wild-type ZJ316; KO, plnJK knockout strain; RE, complemented strain; Over, overexpression strain.

#### 3.4.2 Effect of strain-derived PlnJK on gut microbiota composition

At the genus level, we analyzed the relative abundance of gut microbiota across groups. As shown in Fig.5C and 5D, *Escherichia-Shigella* became dominant in the Ec group, whereas its abundance was significantly lower in the WT, RE, and Over groups compared to the Ec group (p < 0.05, p < 0.01, p < 0.001). Notably, the Over group had significantly lower Escherichia-Shigella abundance than the KO group (p < 0.05). Concurrently, the relative abundances of potentially beneficial genera such as *Parasutterella*, *Clostridium*, and *Bifidobacterium* were increased in the WT and Over groups.

#### 3.4.3 Differential analysis of gut microbiota structure by LEfSe

LEfSe analysis (LDA > 3) was used to identify taxa differentially enriched among the WT, KO, and Over groups. As shown in Fig.5E, the Over group was significantly enriched in *Rikenellaceae*, *Alistipes*, and *Planococcaceae*. The KO group was characterized by enrichment of *Lactobacillaceae* and *Lactobacillus*. The enrichment of *Lactobacillus* in the KO group might be attributed to the exogenously introduced *L. plantarum* ZJ316.

#### 3.4.4 Effect of strain-derived PlnJK on gut short-chain fatty acid levels

SCFAs in fermented samples were quantified by GC-MS. As shown in Fig.5F, multiple SCFAs were significantly elevated in the Over group compared to the Ec group. Specifically, butyrate was significantly higher (p < 0.01), and isovalerate was also markedly increased (p < 0.01) in the Over group. In addition, butyrate, as a primary energy source for colonocytes, plays a key role in maintaining intestinal barrier integrity and anti-inflammatory responses. Collectively, these findings demonstrate that LAB-derived PlnJK promotes the production of beneficial SCFAs by modulating microbial metabolic functions.

## 4 Discussion

This study first evaluated the *in vitro* antibacterial activity and mechanisms of chemically synthesized PlnJK against *E. coli* O104:H4 by determining MIC, membrane permeability, intracellular ATP, and ultrastructural changes. Using an *in vitro* fecal slurry fermentation system, we then assessed its impact on gut microbiota and SCFA metabolism. Furthermore, engineered LAB strains with *plnJK* knockout, complemented expression, and overexpression were constructed and evaluated in the same system for their selective antibacterial and microecological regulatory effects.

The MIC of PlnJK against *E. coli* O104:H4 was 32 μM, confirming its activity against this Gram-negative pathogen. This contrasts with a previous report where PlnJK caused <10% killing of *E. coli* K-12 unless the outer membrane was disrupted(Pal & Srivastava, 2014). K-12 has rough LPS lacking O-antigen due to wbbL inactivation, with a barrier maintained mainly by OmpF/C porins and surface appendages limited to type I fimbriae and flagella(Wang et al., 2021). In contrast, O104:H4 possesses smooth LPS with an intact O104 O-antigen and exhibits a highly complex surface composition: in addition to flagella (H4), it highly expresses aggregative adherence fimbriae AAF/I encoded by the pAA plasmid, multiple long polar fimbriae (Lpf), and outer membrane-anchored SPATEs (Pic, SigA, and SepA)(Bielaszewska et al., 2011; Rasko et al., 2011). Theoretically, smooth LPS should provide a stronger barrier; however, the abundant and highly disordered fimbrial and proteinaceous layers on the O104:H4 surface may serve as anchoring scaffolds for PlnJK, inducing local outer membrane perturbation that enables the peptide to penetrate. K-12 lacks these virulence-associated appendages, and its outer membrane remains a stable barrier. Therefore, the distinct outer membrane surface architecture of O104:H4, comprising smooth LPS with an intact O104 O-antigen and multiple virulence-associated fimbriae and autotransporters, may represent the key structural basis for its susceptibility to PlnJK, consistent with the hypothesis proposed by Rogne et al.(Rogne et al., 2009).

Furthermore, unlike traditional bacteriocins such as nisin, which generally show weak activity against Gram-negative bacteria, PlnJK showed marked anti-Gram-negative activity, attributed to its two-peptide structure and cationic nature that facilitate electrostatic attraction to the negatively charged bacterial surface(Anderssen et al., 1998; Diep et al., 2009; Field et al., 2023). The sustained increase in extracellular conductivity and rapid decline in intracellular ATP indicate that PlnJK disrupts membrane integrity, consistent with Moll et al.(Moll et al., 1999). ATP depletion likely results from dissipation of the proton motive force and direct leakage. SEM and TEM showed plasmolysis and membrane rupture, providing the first ultrastructural evidence of PlnJK disrupting Gram-negative bacterial membranes and offering new insights into cross-genus activity of two-peptide bacteriocins.

In the experiment, *E. coli* O104:H4 infection led to a significant decline in the Shannon index, consistent with reports that EHEC infection disrupts gut microbiota diversity(Croxen et al., 2013). Following the addition of 32 μM chemically synthesized PlnJK, the Chao1 and Shannon indices showed recovery but did not return to the levels of the blank control group. In the LAB-derived PlnJK experiment, the overexpression strain (Over) exhibited a significantly higher Shannon index than the KO and Ec groups, and its Dominance index was significantly lower than that of the KO group. The core reason for this difference lies in the temporal kinetics of effective antimicrobial peptide concentrations: chemically synthesized peptides are prone to proteolytic degradation, adsorption to solid particles, or aggregation in the fecal slurry system, leading to a rapid decline in bactericidal activity over time(Soltani et al., 2021). In contrast, PlnJK-producing LAB—especially the overexpression strain—can continuously secrete the bacteriocin in situ throughout the 24-hour fermentation, maintaining a locally effective concentration, thereby more thoroughly eliminating the pathogen and allowing a time window for the recovery of the commensal microbiota(Mejía-Caballero & Marco, 2026). In the PCoA ordination, although samples from the chemically synthesized PlnJK group were separated from the Ec group, a certain degree of within-group dispersion remained. In contrast, in the LAB-derived PlnJK groups, samples from the WT, RE, and Over groups clustered tightly and were clearly separated from the Ec and KO groups along the first principal coordinate. This indicates that LAB capable of continuously secreting PlnJK can steer the microbiota structure toward a more stable and convergent healthy state. Heilbronner et al. proposed in a review that bacteriocin-mediated microbiome shaping is essentially a form of “niche editing,” the success of which depends on the sustained selective pressure exerted by antimicrobial peptides across both spatial and temporal dimensions(Heilbronner et al., 2021). The strongest community structure normalization effect observed with the overexpression strain in this study aligns well with this concept.

At the genus level, both experiments consistently observed significant inhibition of *Escherichia-Shigella* by PlnJK. In the chemically synthesized PlnJK group, the relative abundance of *Escherichia-Shigella* in the Ec group rose markedly from 26.70% to 36.70%, decreasing to 30.60% after PlnJK addition. In the LAB-derived PlnJK group, the WT, RE, and Over groups all showed significantly lower Escherichia-Shigella abundance compared to the Ec group, with the Over group exhibiting significantly lower abundance than the KO group. This result is consistent with the argument by Sugrue et al. that bacteriocin-mediated “niche clearing” can selectively eliminate competitive pathogens, thereby creating colonization space for beneficial commensals(Sugrue et al., 2024). Notably, the bacterial taxa enriched following both delivery modes displayed pronounced probiotic characteristics. In the chemically synthesized PlnJK group, the relative abundance of *Clostridium* (5.70%) was significantly higher than in the Ctrl (2.50%) and Ec (2.40%) groups. Some species within *Clostridium* (e.g., *Clostridium butyricum* and the *Clostridium leptum* subgroup) are recognized butyrate producers, and their increased abundance aligns with the significant elevation in butyrate levels observed in this study(Guo et al., 2020). In the LAB-derived PlnJK group, the WT and Over groups also showed increased relative abundances of potentially beneficial genera such as *Parasutterella* and *Bifidobacterium*, and LEfSe analysis revealed significant enrichment of *Rikenellaceae* and *Alistipes* in the Over group. Increased abundance of Rikenellaceae has been reported in multiple studies to be associated with a reduced risk of inflammatory bowel disease, while Alistipes has been positively correlated with an anti-inflammatory intestinal environment(De Vos et al., 2022; Wilmanski et al., 2021). These results indicate that the interventional effect of PlnJK extends beyond mere “killing”; rather, it indirectly promotes the ecological recovery of beneficial microbiota through the selective elimination of pathogens.

In this study, GC-MS was employed to quantify seven SCFAs. In the chemically synthesized PlnJK group, compared with the Ec group, the levels of propionate, butyrate, valerate, isovalerate, and hexanoate increased to varying degrees, with significant elevations in isovalerate and butyrate. In the LAB-derived PlnJK group, butyrate was significantly higher in the Over group than in the KO group, isovalerate was also significantly increased, and isobutyrate showed significant upward trends in the Over group. Notably, PlnJK modulated SCFAs in a marked dose-dependent manner: chemically synthesized PlnJK (single addition) produced limited increases, the WT group (basal secretion of the wild-type) had moderate effects, while the Over group (high expression) exhibited the strongest effect. Louis and Flint noted that the abundance of butyrate-producing bacteria in the gut, such as *Clostridium* clusters IV and XIVa, is strongly positively correlated with butyrate production(P. Guo et al., 2020). In this study, the significant proliferation of *Clostridium* (5.70%) in the chemically synthesized PlnJK group and the enrichment of butyrate-producing taxa such as *Rikenellaceae* in the Over group of the LAB-derived PlnJK experiment coincided closely with the concurrent increase in SCFA levels. These findings suggest that PlnJK may eliminate dominant pathogens represented by *Escherichia-Shigella*, thereby releasing occupied niches and nutrient resources, which subsequently allows the expansion of butyrate-producing bacteria and ultimately manifests as elevated SCFA production.

This study has limitations. The fecal slurry model cannot fully replicate the intestinal mucus layer, immune cells, epithelial barrier, and neuro-immune-microbiota axis(Li et al., 2026). Although knockout, complemented, and overexpression strains confirmed *PlnJK*’s specific contribution, the microecological effects of LAB-derived PlnJK may result from synergistic actions of direct bacteriocin killing and intrinsic probiotic effects of the LAB chassis. Future studies could use transcriptomics and metabolomics to dissect PlnJK-driven changes in microbial gene expression and metabolic pathways.

In summary, this study is the first to systematically reveal the antibacterial mechanism of *Lactiplantibacillus plantarum* bacteriocin PlnJK against *E. coli* O104:H4 and its dual modulatory effects on gut microecology. PlnJK disrupts outer membrane integrity, causing ion leakage, ATP depletion, and ultrastructural damage, leading to efficient killing; the selective susceptibility of O104:H4 is linked to its unique outer membrane configuration. Both chemically synthesized and LAB-derived PlnJK selectively eliminate Escherichia-Shigella, indirectly preserving microbial diversity, enriching butyrate-producing bacteria, and upregulating SCFA metabolism. Importantly, LAB-mediated sustained in situ delivery outperformed chemical synthesis in restoring diversity, reshaping community structure, and enhancing metabolic function. These findings provide a new structural biology perspective on cross-genus antibacterial mechanisms of two-peptide bacteriocins and support the design of anti-EHEC live biotherapeutic products based on LAB-derived bacteriocins. Future studies should use transcriptomics and metabolomics to further dissect PlnJK-driven microbial gene expression and metabolic remodeling and validate in vivo efficacy and safety in animal infection models.

## Author contributions

**Shumin Wang:** Conceptualization, Formal analysis, Writing - original draft. **Jiaqian Yan:** Methodology, Data curation, Validation. **Bin Yu:** Formal analysis, Visualization, Writing – original draft. **Ning Ding:** Investigation, Project administration. **Xiaolu Li:** Project administration, Visualization. **Shuxin Wang:** Supervision**. Jiahao Jiang:** Investigation. **Xiaodan Zhao:** Project administration. **Qing Gu:** Resources, Conceptualization, Supervision. **Ping Li:** Funding acquisition, Supervision, Project administration, Writing – review & editing.

## Conflicts of interest

The authors declare no competing interests.

## Data availability

The data supporting the findings of this study are available within the article.

## Acknowledgements

This work was supported by the National Natural Science Foundation of China [No. 32272301].

